# Alpha-Asarone Attenuates Alcohol-Induced Hepatotoxicity in a Murine Model by Ameliorating Oxidative Stress, Inflammation, and Modulating Apoptotic-Autophagic Cell Death

**DOI:** 10.1101/2023.10.24.563724

**Authors:** Amir Mohamed Abdelhamid, Nada A.M. Ali, Norhan M. El Sayed, Asmaa Radwan

## Abstract

Alcoholic liver disease (ALD) is a major cause of chronic liver injury characterized by steatosis, inflammation, and fibrosis. This study explored the hepatoprotective mechanisms of alpha-asarone in a mouse model of chronic-binge alcohol feeding. Adult male mice were randomized into control, alcohol, and alcohol plus alpha-asarone groups. Serum aminotransferases and histopathology assessed liver injury. Oxidative stress was evaluated via malondialdehyde content, glutathione, superoxide dismutase, and catalase activities. Pro-inflammatory cytokines TNF-α, IL-1β, and IL-6 were quantified by ELISA. P53-mediated apoptosis was determined by immunohistochemistry. Key autophagy markers AMPK, Beclin-1, and LC3 were examined by immunoblotting. Alcohol administration increased serum ALT, AST and ALP, indicating hepatocellular damage. This liver dysfunction was associated with increased oxidative stress, inflammation, p53 expression and altered autophagy. Alpha-asarone treatment significantly decreased ALT, AST and ALP levels and improved histological architecture versus alcohol alone. Alpha-asarone also mitigated oxidative stress, reduced TNF-α, IL-1β and IL-6 levels, ameliorated p53 overexpression and favorably modulated autophagy markers. Our findings demonstrate that alpha-asarone confers protective effects against ALD by enhancing antioxidant defenses, suppressing hepatic inflammation, regulating apoptotic signaling, and restoring autophagic flux. This preclinical study provides compelling evidence for the therapeutic potential of alpha-asarone in attenuating alcohol-induced liver injury and warrants further evaluation as a pharmacotherapy for ALD.

## 1. Introduction

Alcohol abuse poses a major hazard to people’s health. Alcohol misuse is the top cause of death among people between the ages of 15 and 49 and the eighth largest risk factor for death and disability-adjusted life years globally [1]. Numerous morphological and functional alterations brought on by excessive alcohol use might result in liver damage [2]. Liver injury and disease can be triggered by a diverse set of metabolic, toxic, and inflammatory insults. Alcohol is metabolized by an oxidative metabolic pathway that takes place in the endoplasmic reticulum of hepatocytes [3]. With the help of NADPH and oxygen, CYP2E1 converts ethanol to acetaldehyde and then acetaldehyde to acetate. When ethanol is converted into acetaldehyde, reactive oxygen species (ROS) are formed, which contribute to alcohol toxicity [3]. Alcohol always leads to the activation of apoptosis, necrosis, and autophagy in liver cells [4]. The mitochondrial apoptotic pathway, or intrinsic pathway, is implicated in many death scenarios, such as DNA damage or hormone or growth factor deficiencies. The death receptor pathway, or extrinsic pathway, is mainly initiated by the binding of death receptor ligands to death receptors [5].

The cornerstone of treating people with ALD is achieving and maintaining alcohol abstinence because the effectiveness of ALD treatment is hampered by patients who continue to drink [6]. Corticosteroids are currently used in patients with alcoholic hepatitis. The short-term survival benefit of corticosteroid treatment has been confirmed in a recent meta-analysis that included corticosteroids (e.g., methylprednisolone) or TNF-inhibitor therapy (e.g., pentoxifylline) for alcoholic hepatitis [7–9]. However, these treatments yield conflicting outcomes and are associated with increased rates of infection [10]. Therefore, it’s important to focus on new drugs that have fewer limitations and improve patient survival.

Studies have suggested that both alpha and beta asarone may be helpful in the treatment of neurodegenerative illnesses, including Alzheimer’s and Parkinson’s disease, considering their neuroprotective properties, anti-apoptotic effects against neuronal apoptosis, and attenuation of neuroinflammation. Additionally, based on the now available evidence, alpha-asarone may have good therapeutic potential for the treatment of CNS disorders (depression, anxiety, drug dependence, pain), CVS disorders (hyperlipidemia, coronary artery diseases), and cholestasis [11]. Numerous studies in vitro and in vivo on alpha-asarone have proven that it has antioxidant action [12]. This research sought to examine how alpha-asarone may affect alcohol-induced liver injury.

## 2. Materials and Methods

### 2.1. Animals

Forty male Swiss albino mice weighing 30 ± 3 g were obtained from the animal house of the faculty of pharmacy at Suez Canal University (Ismailia, Egypt). Mice were housed in a temperature-controlled environment (22 ± 3 ◦C), 45%–55% humidity, and a 12-h light/dark cycle. After baseline behavioral assessment, the animals were allowed to acclimatize for two weeks before the experiment. Throughout the experiment, mice were weighed and regularly monitored for any indications of distress. The experiment protocols were performed in compliance with the Animal Ethics Committee’s guidelines at the Faculty of Pharmacy, Suez Canal University (License number 202204MA2), in accordance with the Canadian Council on Animal Care Guidelines.

### 2.2. Drugs and chemicals

Alpha-asarone powder was obtained from ThermoFisher Gmbh (Kandel, Germany) and was prepared as a 0.1% w/v suspension in 0.9% sodium chloride. Alcohol was purchased from Alamia Chemicals Company (Sharqia, Egypt) as 96% (vol/vol) ethanol (ETOH). The Lieber-DeCarli control liquid diet (product No. F1259SP), Lieber-DeCarli ethanol liquid diet (product No. F1258SP), and maltose dextrin (product No. 3653) were obtained from Bio-Serv (NJ, USA).

### 2.3. Experimental design

The NIAAA model (chronic ethanol feeding plus a single binge), which is easy to use, shows a clear rise in ALT and steatosis, is short-term feeding with no mortality rate, and does not cause liver fibrosis, was employed in the study to create the current study model [13]. Mice were divided into five groups (Figure 1): Group I (Normal): received control liquid diet; Group II (ALC): received ethanol liquid diet; Group III (ALC+ASR 5): received ethanol liquid diet and alpha-asarone 5 mg/kg/day; Group IV (ALC+ASR 10): received ethanol liquid diet and alpha-asarone 10 mg/kg/day; Group V (ASR 10): received control liquid diet and alpha-asarone 10 mg/kg/day. On day 1, mice were acclimatised to the Lieber-DeCarli control liquid diet (0% EtOH) as the only source of water and food, and treatments were administered intraperitoneally once a day beginning on day 2. Alcoholic groups were fed an ethanol-containing liquid diet, with a gradual increase in EtOH concentrations from 1% to 4% (vol/vol) from day 2 to day 5. Then, beginning on day 6 and for 10 days, feeding tubes were filled with a 5% (vol/vol) ethanol-containing liquid diet [14]. On the last day, the gavage volume of 31.5% (vol/vol) ethanol solution was administered for ethanol liquid diet groups, while the gavage volume of 45.0% (wt/vol) maltose dextrin solution was administered for control liquid diet groups. The mice were scarified 9 hours after the gavage [12]. Blood samples were collected before the sacrifice from the orbital sinus. After sacrifice, livers were isolated and divided into three parts. The first part was prepared as 10% homogenate for assessment of biochemical parameters; the second one was prepared for histopathological and immunohistochemical examinations; and the third one was preserved at -80°C for further analyses.

**Figure 1:**
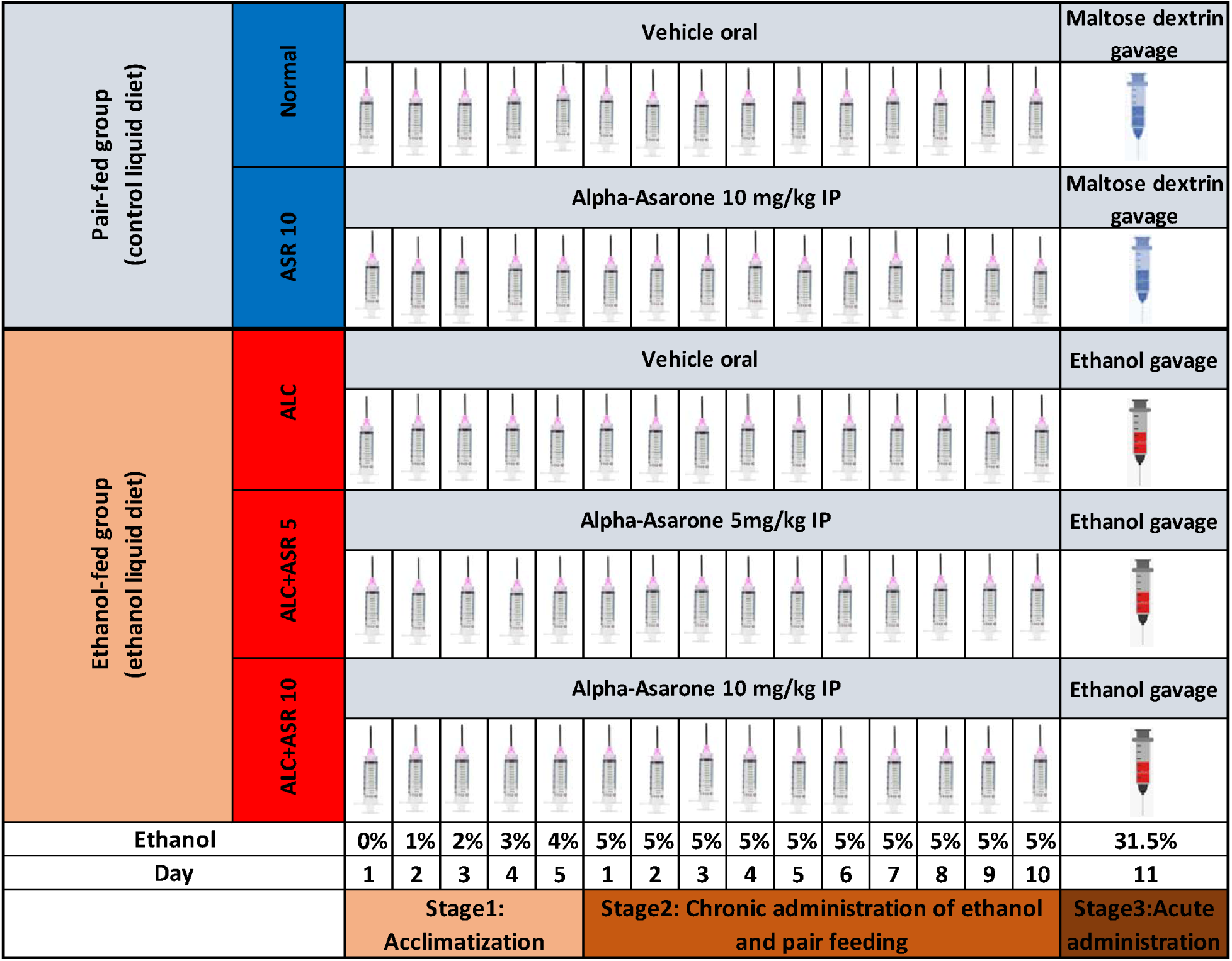
Schematic representation of the experimental design.

### 2.4. Assessment of liver enzymes

Using a kit provided by TECO Diagnostics (CA, USA), the serum aspartate aminotransferase (AST) and alanine aminotransferase (ALT) levels were determined [15]. The QuantiChromTM Alkaline Phosphatase Assay Kit (DALP-250), provided by Bioassay Systems (CA, USA), was used to test alkaline phosphatase (ALP).

### 2.5. Quantitative determination of the levels of inflammatory cytokines

A 10% liver homogenate was prepared using phosphate-buffered saline. The levels of interleukin-1 (IL-1), interleukin-6 (IL-6), and tumor necrosis factor-α (TNF-α) were measured by ELISA technique using ELISA kits from ShangHai BlueGene Biotech CO, LTD (IL-1, Cat. No. MBS825017; IL-6, Cat. No. MBS355410; TNF-, Cat. No. MBS2507393) following the manufacturer’s instructions.

### 2.6. Assessment of antioxidant enzymes

Reduced glutathione (GSH) concentration, superoxide dismutase (SOD) activity, catalase (CAT) activity, and malondialdehyde (MDA) content were measured using ELISA kits from ShangHai BlueGene Biotech CO, LTD (SOD, Cat. No. MBS036924; GSH, Cat. No. E02G0367; CAT, Cat. No. MBS006963; MDA, Cat. No. LS-F28018) following the manufacturer’s instructions.

### 2.7. Assessment of autophagy biomarkers

Protein expression levels of AMPK, Beclin-1, and LC3 were assessed utilizing the western blotting technique [16]. TriFast (Peqlab, VWR company) was employed for simultaneous isolation of RNA, DNA, and protein. Anti-AMPK alpha 1 antibody (2B7), anti-Beclin 1 antibody (EPR19662), and anti-LC3 antibody (ab36817), all sourced from Abcam, USA, were utilized for this analysis.

### 2.8. Histopathological examination of liver tissue and immunohistochemical analysis

Following fixation with 10% formalin, the samples were embedded in paraffin. Subsequently, 3 µm thick sections were obtained from each paraffin block, mounted on glass slides, and subjected to hematoxylin and eosin (H&E) staining. The evaluation of these stained sections was performed by an independent pathologist [17]. Slide scanning and image processing were conducted using the ImageJ scanner and viewer software (LOCI, University of Wisconsin, US). The liver tissue sections were assessed based on Ishak’s modified histological activity index (HAI) grading system. The identified conditions included periportal or periseptal interface hepatitis (piecemeal necrosis) graded from 0 to 4, confluent necrosis graded from 0 to 6, focal (spotty) lytic necrosis, apoptosis, and focal inflammation graded from 0 to 4, and portal inflammation graded from 0 to 4 [18].

For immunohistochemistry (IHC) staining, 4-micrometer thick sections were obtained from selected paraffin blocks. Slides were incubated with primary anti-P53 antibodies (53kDa) obtained from ABclonal (Woburn, USA). Subsequently, the appropriate secondary antibody was applied. A faint counterstaining with hematoxylin for 30 seconds was performed prior to dehydration and mounting. Positive cells exhibiting a nuclear and/or cytoplasmic response to p53 were identified. The modified Allred scoring method was employed to conduct a semi-quantitative analysis of the stained tissue sections [19]. The quantification of positive cells involved counting them in three distinct high-power fields (hpf) at a magnification of 400x. The average number of positive cells was determined. The final grades were obtained by combining the scores for the percentage of positive cells (ranging from 0 to 5) and the staining intensity of the cytoplasm (ranging from 0 to 3).

### 2.9. Statistical analysis

Parametric data are expressed as mean ± standard deviation (S.D). Statistical analysis of the results was performed using GraphPad Prism (version 9.5.0) software. One-way analysis of variance (ANOVA) followed by Tukey’s multiple comparison test was employed for analyzing parametric data. Histopathological scores are presented as the median with interquartile range, and their comparison was conducted using Kruskal-Wallis followed by Dunn’s multiple comparison test. Statistical significance was considered at a threshold of p < 0.05.

## 3. Results

### 3.1 Effect of alpha-asarone on AST, ALT, ALP in mice with alcohol-induced liver injury

In the current study, alcohol was used to induce alcoholic liver injury. To prove that alpha-asarone ameliorates alcoholic liver dysfunction, liver enzymes ALT, AST, and ALP were measured. Results showed that the ALC group fed with an ethanol liquid diet had elevated levels of serum ALT, AST, and ALP compared with the normal group. Treatment with alpha-asarone in the ALC+ASR 5 and ALC+ASR 10 groups showed a significant decrease in liver enzymes compared with the ALC group (Table 1).

**Table 1.** Effect of alpha-asarone on Aspartate Aminotransferase (AST), Alanine Aminotransferase (ALT) and Alkaline Phosphatase (ALP) in mice with alcoholic liver injury.

|  | Normal | ALC | ALC+ASR 5 | ALC+ASR 10 | ASR 10 |
| --- | --- | --- | --- | --- | --- |
| **AST** | 44.64 ±<br>3.189 | 144.5 ± 6.12* | 90.76 ± 3.99*# | 59.38 ± 3.66*#\$ | 40.14 ±<br>1.258# |
| **ALT** | 29.39±1.605 | 89.89±2.266* | 62.01±2.859*# | 45.63±2.504*#\$ | 26.38±1.685# |
| **ALP** | 70.88±4.357 | 211.4±7.914* | 118.0±2.958*# | 99.26±2.846*#\$ | 58.26±2.846*# |

Values are presented as mean ± SD. Data were compared using one-way analysis of variance (ANOVA) followed by Tukey's multiple comparison test at $P < 0.05$ . \*, significantly different from the normal group. #, significantly different from the ALC group. \$, significantly different from ALC+ASR 5 group.

### 3.2. Effect of alpha-asarone on liver morphology in mice with alcohol-induced liver injury

Histopathological examination of liver tissues stained with hematoxylin and eosin (H&E) (Figure 2) revealed distinct observations among the experimental groups. In the normal group, liver tissue exhibited uniformity with hepatocytes arranged in regular plates, devoid of any signs of injury. In contrast, the ALC group displayed multiple regions of portal inflammation, along with evidence of interface hepatitis, areas of confluent necrosis, and scattered foci of lobular inflammation, as indicated by a higher histopathological score compared to the normal group (Figure 2A). Upon treatment with alpha-asarone in the ALC+ASR 5 group, mild expansion of portal tracts accompanied by inflammation was observed, along with a few scattered foci of lobular inflammation and notable steatosis in hepatocytes. In the ALC+ASR 10 group, treatment with alpha-asarone resulted in mild portal tract inflammation limited to a few portal tracts, along with evidence of mild interface hepatitis, absence of confluent necrosis or foci of lobular inflammation, and no steatosis in hepatocytes. These observations led to a significant decrease in the histopathological score compared to the ALC group. Lastly, the ASR 10 group exhibited multiple areas of portal inflammation with mild interface hepatitis and scattered foci of lobular inflammation.

**Figure 2:**
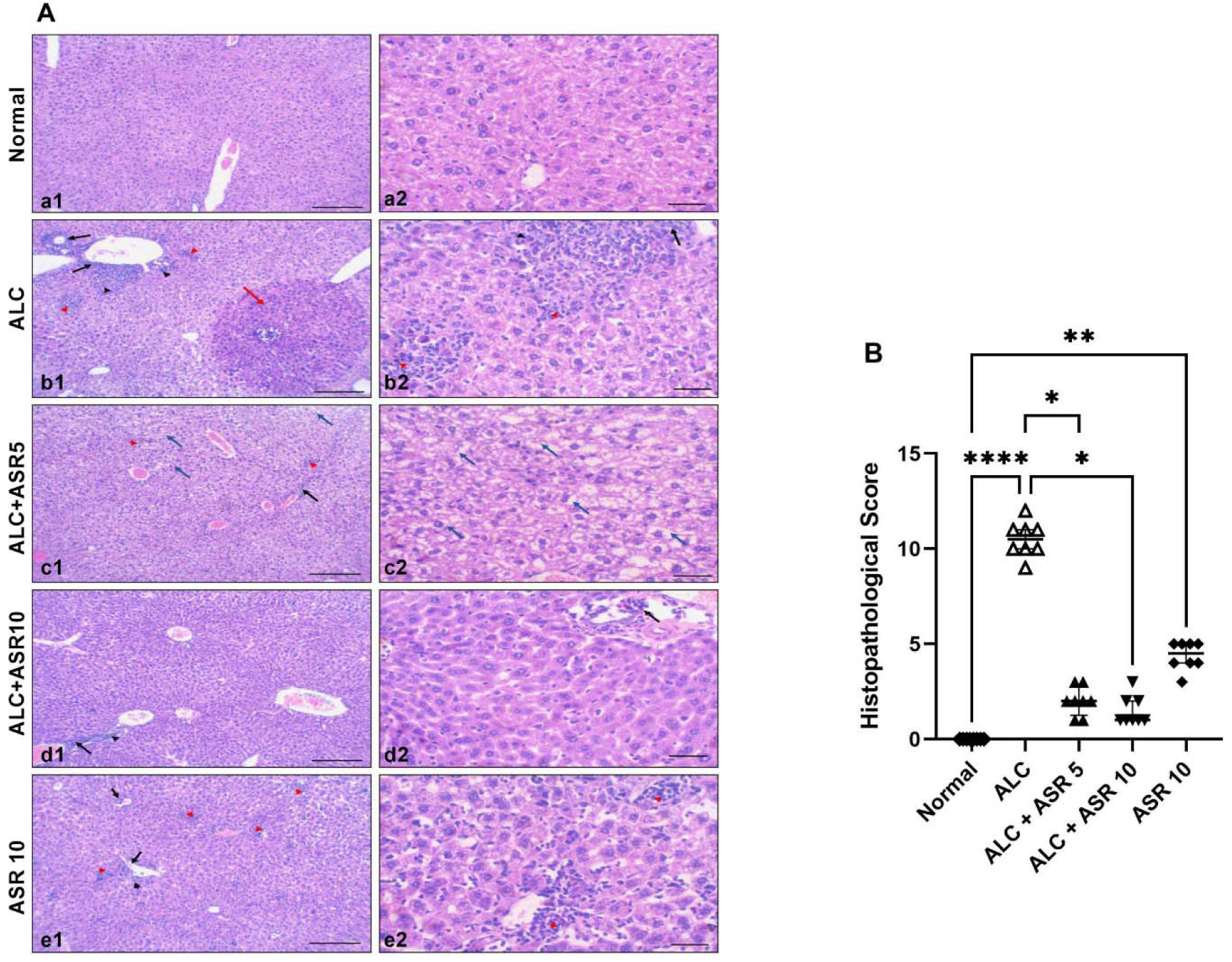
Impact of alpha-asarone on liver morphology in mice with alcoholic liver injury. (A) Microscopic pictures of H&E-stained liver sections showed uniform liver tissue with plates of uniform hepatocytes with no evidence of injury in the normal group (a1, a2). Multiple areas of portal inflammation (black arrows) with evidence of interface hepatitis (black arrowheads), an area of confluent necrosis (red arrow), and scattered foci of lobular inflammation (red arrowheads) in the ALC group (b1, b2). Mild expansion of portal tracts with inflammation (black arrows), few scattered foci of lobular inflammation (red arrowheads), and significant steatosis in hepatocytes (blue arrows) in the ALC+ASR 5 group (c1, c2). Mild portal tract inflammation in a few portal tracts (black arrow) with evidence of interface hepatitis (black arrowheads), no confluent necrosis or foci of lobular inflammation, and no steatosis in hepatocytes in the ALC+ASR 10 group (d1, d2). Multiple areas of portal inflammation (black arrows) with evidence of mild interface hepatitis (black arrowheads) and scattered foci of lobular inflammation (red arrowheads) in the ASR 10 group (e1, e2). 1 (x:10, bar 100); 2 (x:40, bar 20). (B) Effect of alpha-asarone on mice with alcoholic liver disease’s histological scores. The data is displayed as mean ± SD (n □ 8). * p < 0.05, ** p < 0.01, **** p < 0.0001. The normal group received a control liquid diet; the ALC group received an ethanol liquid diet; the ALC+ASR 5 group received an ethanol liquid diet and alpha-asarone 5 mg/kg; the ALC+ASR 10 group received an ethanol liquid diet and alpha-asarone 10 mg/kg; and the ASR 10 group received a control liquid diet and alpha-asarone 10 mg/kg.

### 3.3. Effect of alpha-asarone on IL-1, IL-6, and TNF-**α** in mice with alcohol-induced liver injury

The impact of alpha-asarone on inflammatory markers in the liver, specifically IL-1, IL-6, and TNF-α, is depicted in Figure 3A, 3B, and 3C, respectively. In the ALC group, there was a significant elevation in the levels of IL-1, IL-6, and TNF-α (50.53 ± 2.319 pg/g tissue, p < .0001; 40.18 ± 1.737 pg/g tissue, p < .0001; and 56.76 ± 3.237 pg/g tissue, p < .0001, respectively) when compared to the normal group. Conversely, treatment with alpha-asarone in the ALC+ASR 5 group significantly mitigated these increases in comparison to the ALC group (IL-1: 30.62 ± 1.424 pg/g tissue, p < .0001; IL-6: 25.44 ± 1.319 pg/g tissue, p < .0001; and TNF-α: 32.50 ± 1.789 pg/g tissue, p < .0001). Additionally, treatment in the ALC+ASR 10 group demonstrated a substantial reduction in the levels of IL-1, IL-6, and TNF-α (23.69 ± 1.870 pg/g tissue, p < .0001; 17.09 ± 1.210 pg/g tissue, p < .0001; and 23.04 ± 1.580 pg/g tissue, p < .0001, respectively) when compared to the ALC group. Also, the ASR 10 group showed a mildly significant decrease in the levels of inflammatory markers (IL-1: 9.763 ± 0.9471 pg/g tissue, p < .0001; IL-6: 8.225 ± 1.002 pg/g tissue, p < .0001; and TNF-α: 10.33 ± 0.6386 pg/g tissue, p < .0001) when compared to the normal group. Finally, treatment with a higher dose of alpha-asarone in the ALC+ASR 10 group showed a higher amelioration in inflammatory cytokines compared to the ALC+ASR 5 group.

**Figure 3:**
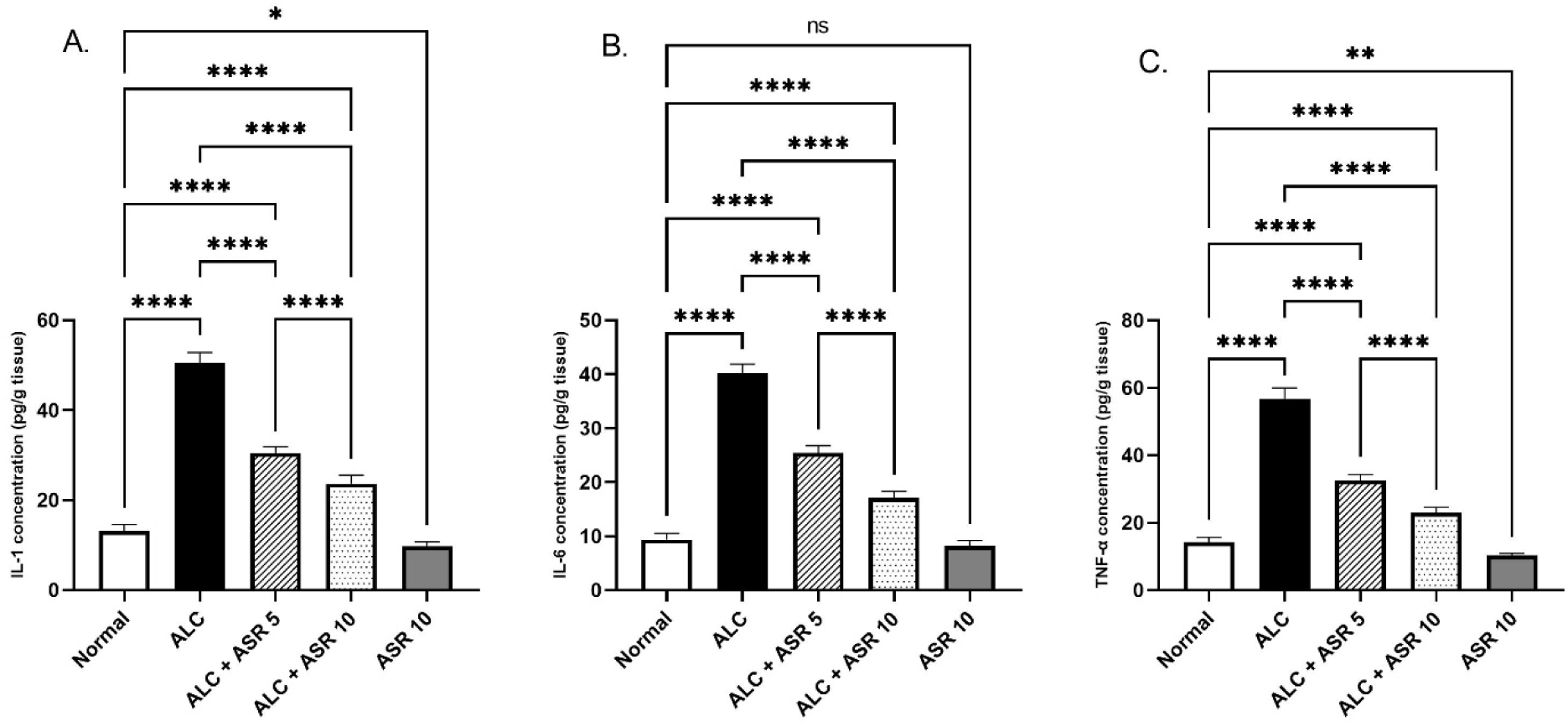
Effect of alpha-asarone on IL-1 (A), IL-6 (B), and TNF-α (C) in the livers of mice with alcoholic liver injury. Data is displayed as the mean ± SD (n□8). Ns, non-significant, * p < 0.05, ** p < 0.01, **** p < 0.0001. The normal group received a control liquid diet; the ALC group received an ethanol liquid diet; the ALC+ASR 5 group received an ethanol liquid diet and alpha-asarone 5 mg/kg; the ALC+ASR 10 group received an ethanol liquid diet and alpha-asarone 10 mg/kg; and the ASR 10 group received a control liquid diet and alpha-asarone 10 mg/kg.

### 3.4. Effect of alpha-asarone on GSH, SOD, CAT, and MDA in mice with alcohol-induced liver injury

The influence of alpha-asarone on various parameters, including GSH concentration (Figure 4A), SOD activity (Figure 4B), CAT activity (Figure 4C), and MDA content in the liver (Figure 4D), was investigated. In the ALC group, there was a reduction in GSH concentration (14.36 ± 1.307 Mmol/g tissue, p < .0001), decreased SOD activity (10.36 ± 0.8245 U/g tissue, p < .0001), and a decline in CAT activity (11.89 ± 0.9598 U/g tissue, p < .0001) compared to the normal group. Conversely, the ALC group exhibited an increase in MDA content (42.48 ± 1.917 ng/g tissue, p < .0001) compared to the normal group. Treatment with alpha-asarone in the ALC+ASR 5 group resulted in a significant elevation in GSH concentration (30.48 ± 1.593 Mmol/g tissue, p < .0001), increased SOD activity (25.05 ± 1.023 U/g tissue, p < .0001), and enhanced CAT activity (24.94 ± 1.327 U/g tissue, p < .0001) compared to the ALC group. Additionally, treatment in the ALC+ASR 5 group showed a reduction in MDA content (24.55 ± 1.204 ng/g tissue, p < .0001) compared to the ALC group. Moreover, treatment with alpha-asarone in the ALC+ASR 10 group demonstrated a significant elevation in GSH concentration (41.01 ± 1.743 Mmol/g tissue, p < .0001), increased SOD activity (31.75 ± 1.757 U/g tissue, p < .0001), and enhanced CAT activity (33.71 ± 1.736 U/g tissue, p < .0001) compared to the ALC group. Furthermore, treatment in the ALC+ASR 10 group exhibited a reduction in MDA content (18.90 ± 1.350 ng/g tissue, p < .0001) compared to the ALC group. Also, In ASR 10, there was also a little increase in SOD activity (48.49 1.972 U/g tissue, p.0001) and an increase in CAT activity (57.84 1.530 U/g tissue, p.0001) compared to the control group. Finally, the ALC+ASR 10 group received a greater dose of alpha-asarone, which resulted in a substantial rise in GSH concentration, improved SOD activity, and enhanced CAT activity compared to the ALC+ASR 5 group. Furthermore, therapy in the ALC+ASR 10 group reduced MDA content as compared to the ALC+ASR 5 group.

**Figure 4:**
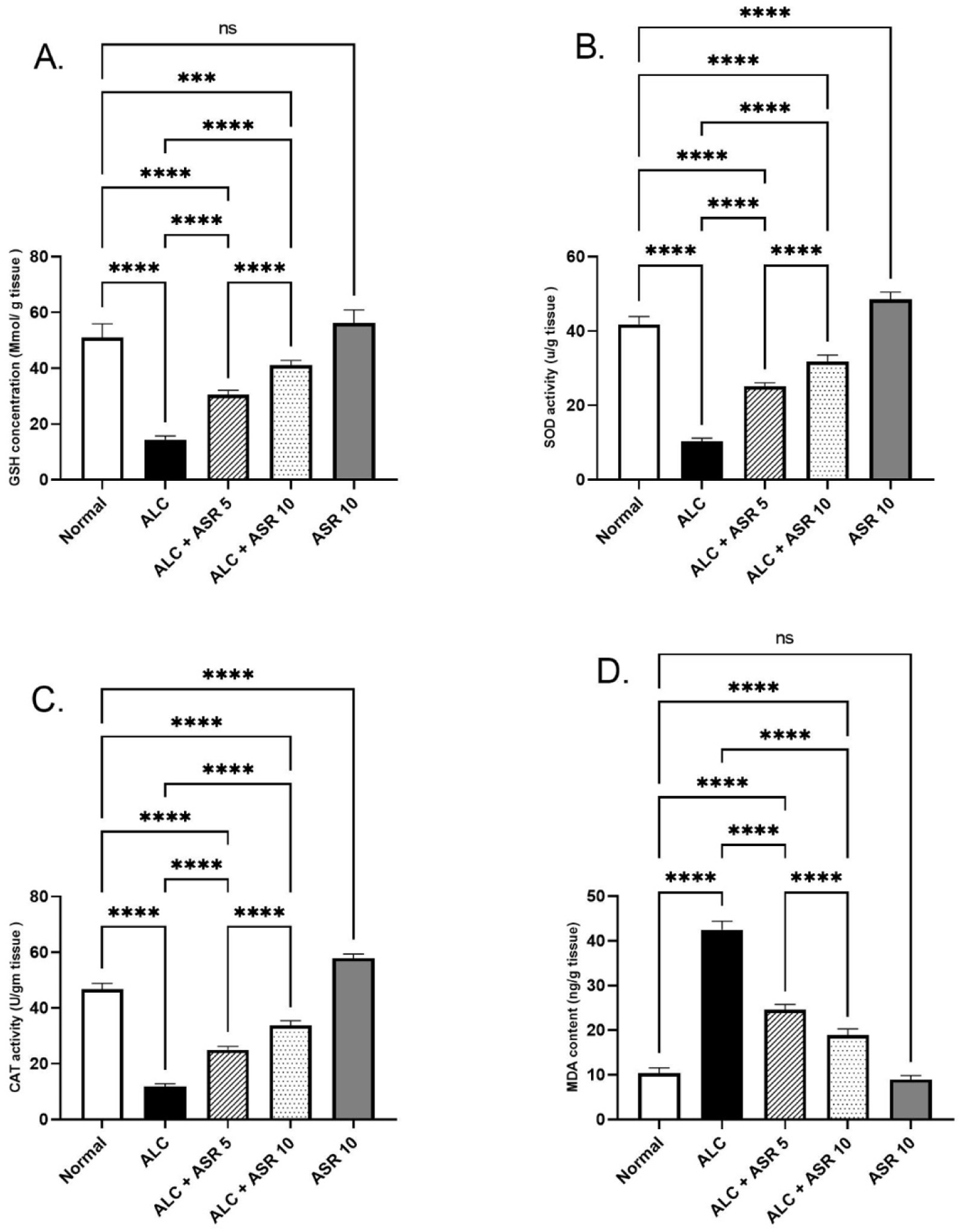
Effect of alpha-asarone on the GSH concentration (A); SOD activity (B); CAT activity (C); and MDA content (D) in the livers of mice with alcoholic liver injury. Data is displayed as the mean ± SD (n□8). Ns, non-significant, *** P<0.001, ****p<0.0001. The normal group received a control liquid diet; the ALC group received an ethanol liquid diet; the ALC+ASR 5 group received an ethanol liquid diet and alpha-asarone 5 mg/kg; the ALC+ASR 10 group received an ethanol liquid diet and alpha-asarone 10 mg/kg; and the ASR 10 group received a control liquid diet and alpha-asarone 10 mg/kg.

### 3.5. Effect of alpha-asarone on AMPK, Beclin-1, and LC3 in mice with alcohol-induced liver injury

The impact of alpha-asarone on the protein expression levels of AMPK, Beclin-1, and LC3, serving as indicators of autophagy, in the liver is depicted in Figure 5A, 5B, 5C and 5D, respectively. In comparison to the normal group, the ALC group exhibited a notable decrease in AMPK protein levels, while the protein levels of Beclin-1 and LC3 were significantly elevated. Conversely, treatment with alpha-asarone in the ALC+ASR 10 group effectively attenuated the decline in AMPK protein levels and demonstrated a significant reduction in Beclin-1 and LC3 protein levels compared to the ALC group. It is worth noting that treatment with alpha-asarone in the ALC+ASR 5 group showed no significant change in these protein expression.

**Figure 5:**
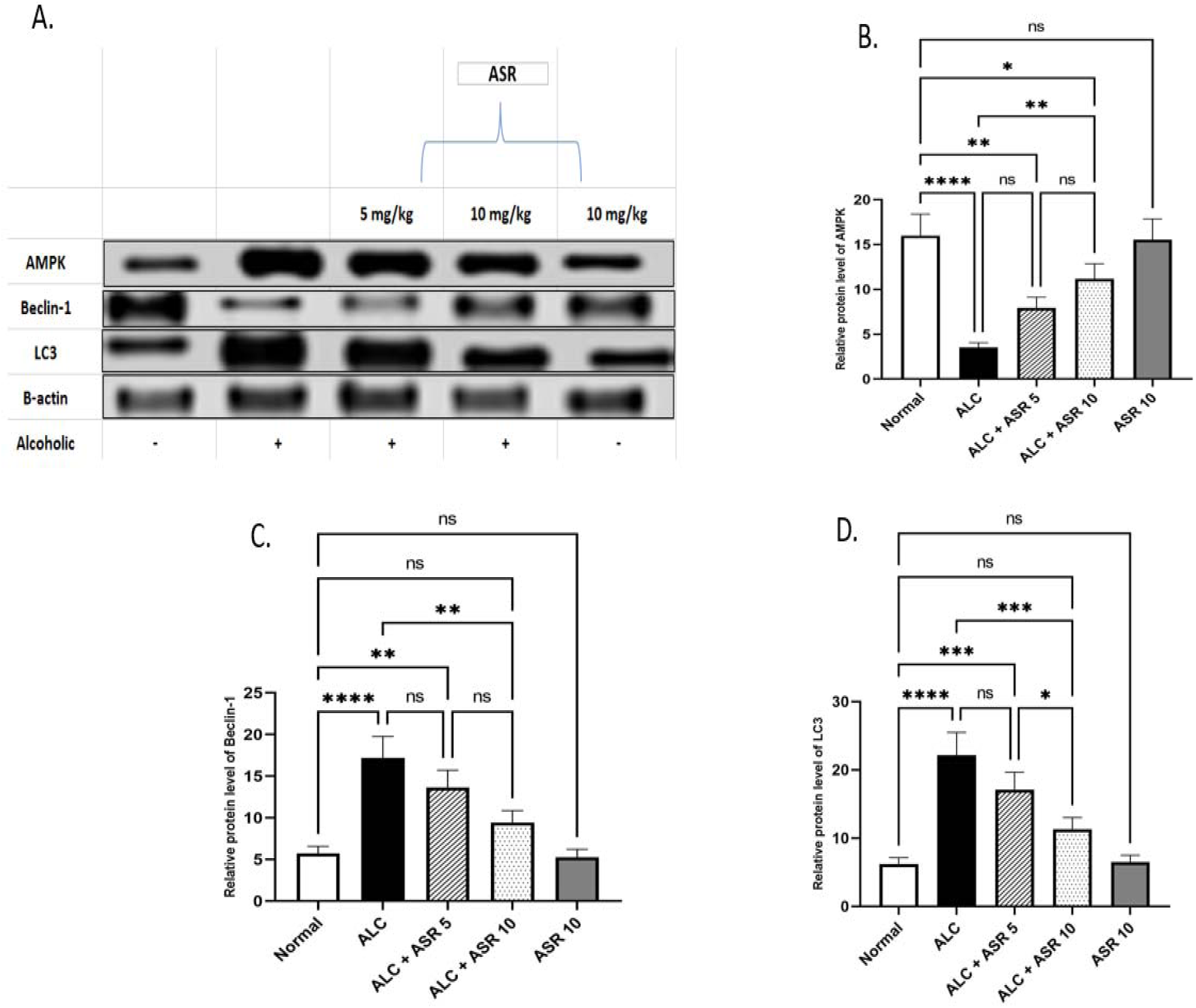
Effect of alpha-asarone on autophagy markers in the livers of mice with alcoholic liver injury. (A) Western blotting technique used in the measurement of the autophagy markers. (B) Relative protein level of AMPK. (C) Relative protein level of Beclin-1. (D) Relative protein level of LC3. The data is displayed as the mean ± SD (n□8). Ns, non-significant, *p<0.05, **p<0.01, *** P<0.001, ****p<0.0001. The normal group received a control liquid diet; the ALC group received an ethanol liquid diet; the ALC+ASR 5 group received an ethanol liquid diet and alpha-asarone 5 mg/kg; the ALC+ASR 10 group received an ethanol liquid diet and alpha-asarone 10 mg/kg; and the ASR 10 group received a control liquid diet and alpha-asarone 10 mg/kg.

### 3.6. Effect of alpha-asarone on the expression of P53 in mice with alcohol-induced liver injury

The impact of alpha-asarone on p53, serving as an apoptosis marker (Figure 6), was assessed through immunohistochemical analysis using an anti-P53 antibody. The normal group exhibited weak scattered positivity for P53 in the nuclei of hepatocytes. In contrast, the ALC group demonstrated strong nuclear and cytoplasmic expression of P53 in hepatocytes. However, treatment with alpha-asarone in the ALC+ASR 5 and ALC+ASR 10 groups significantly reduced the expression of P53 in the cytoplasm of hepatocytes. Statistical analysis further confirmed that the normal group had a low percentage of stained areas, while the ALC group showed an increase in the stained area. Moreover, the groups treated with alpha-asarone exhibited a marked decrease in the stained area compared to the ALC group.

**Figure 6.**
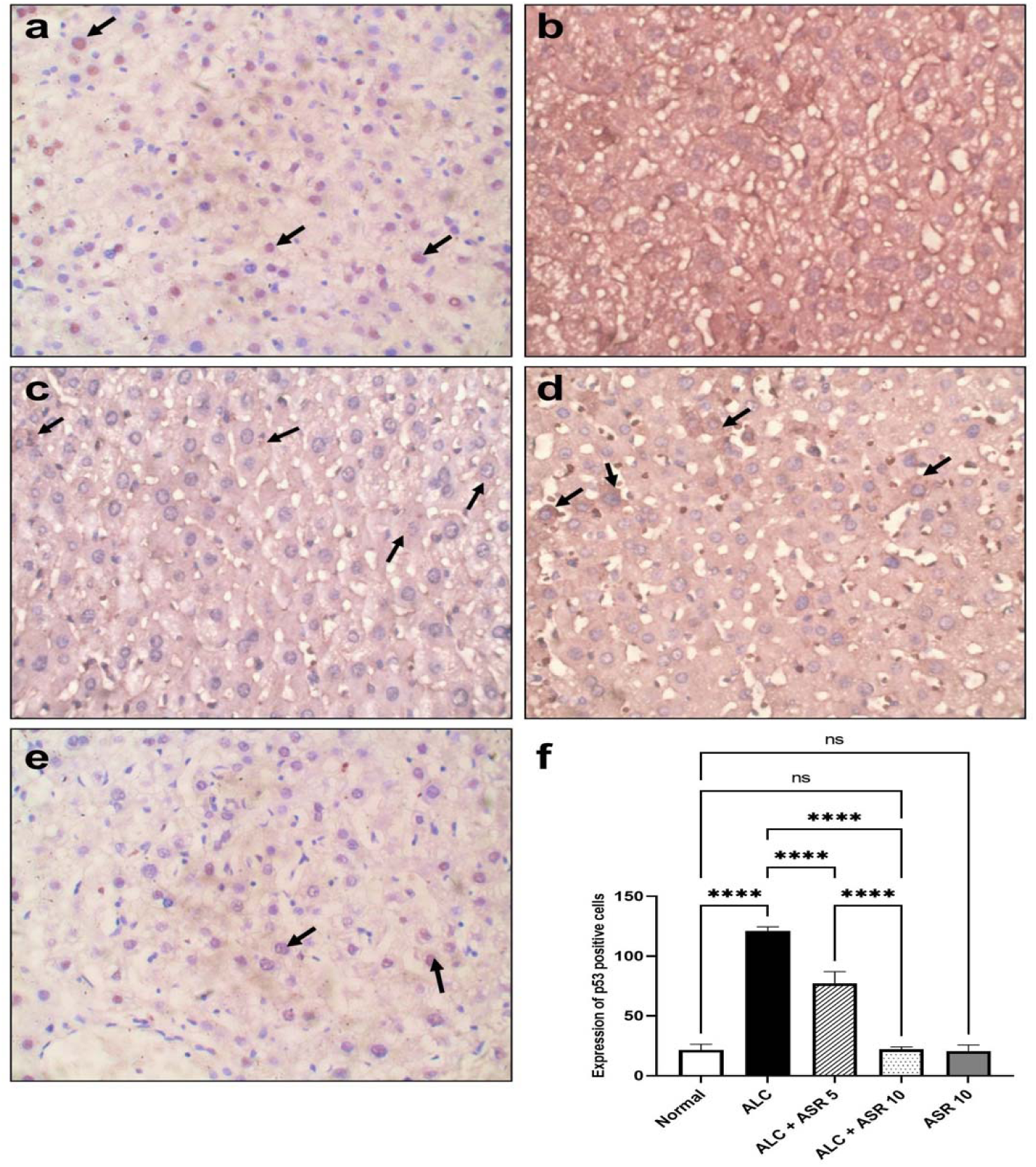
The effect of alpha-asarone on p53 expression in the livers of mice with alcoholic liver injury. The hepatocyte nuclei of the normal group (a) had weakly dispersed positives for P53 (black arrows). P53 was strongly expressed in the nucleus and cytoplasm of hepatocytes from the ALC group (b). In the treated groups ALC+ASR 5 and ALC+ASR 10, there was a considerable decrease in the intensity of P53 expression in the cytoplasm of hepatocytes (black arrows) (c, d)). (f) Effect of alpha-asarone on the immunohistochemical score of p53 positive cells in mice with alcoholic liver injury. The data is displayed as the mean ± SD (n = 8). Ns, non-significant, ****p<0.0001. The normal group received a control liquid diet; the ALC group received an ethanol liquid diet; the ALC+ASR 5 group received an ethanol liquid diet and alpha-asarone 5 mg/kg; the ALC+ASR 10 group received an ethanol liquid diet and alpha-asarone 10 mg/kg; and the ASR 10 group received a control liquid diet and alpha-asarone 10 mg/kg.

## 4. Discussion

Alcohol consumption and related disorders impose a substantial economic burden, resulting in increased healthcare costs and societal consequences. Notably, individuals suffering from alcoholic liver disease (ALD) generate direct healthcare expenses for medical treatment, in addition to indirect costs associated with decreased workforce productivity [20]. The interrelation between the average volume of drinking, pattern of drinking, and the development of diseases is mediated through three key intermediate pathways: intoxication, dependency, and the complex interplay of harmful and beneficial biological effects of alcohol on various organs and tissues [21]. Hepatic steatosis, alcoholic hepatitis, cirrhosis, and liver cancer can all result from long-term heavy drinking [22]. Alcoholic cirrhosis can advance to decompensation and liver cancer despite abstinence, in contrast to hepatic steatosis and, possibly, fibrosis, which may be reversible with the cessation of alcohol usage [23]. Therefore, developing new drugs either for the prevention or treatment of alcoholic liver disease is urgently needed.

Previous studies have suggested potential therapeutic applications of alpha-asarone in treating cholestasis, cardiovascular system (CVS) disorders, and central nervous system (CNS) diseases [11]. Furthermore, extensive in vitro and in vivo investigations have demonstrated the antioxidant properties of alpha-asarone [12]. Given these findings, the present study aimed to investigate the potential of alpha-asarone in the treatment of alcoholic liver disease (ALD).To induce ALD, the NIAAA model, which involves chronic ethanol feeding along with a single binge, was employed [13].

ALD is often characterized by a lack of noticeable symptoms in its early stages and even in patients with compensated cirrhosis, making diagnosis reliant on clinical suspicion, laboratory tests, and invasive or noninvasive techniques [24]. Elevated liver enzymes, such as alanine aminotransferase (ALT), aspartate aminotransferase (AST), and alkaline phosphatase (ALP), are commonly observed in patients with ALD due to the direct toxic effects of alcohol on the liver [25]. In our study, the ALC group exhibited a marked increase in serum levels of ALT, AST, and ALP, indicating liver injury. However, treatment with alpha-asarone (5, 10 mg/kg) mitigated these elevations in enzyme markers, which indicates that alpha-asarone has a curative effect on ALD, aligning with the findings of Tathe et al., who demonstrated the hepatoprotective effects of alpha-asarone in rats [26]. Their study also demonstrated a reduction in liver enzyme markers, supporting the notion that alpha-asarone holds promise as a therapeutic intervention for liver diseases.

Furthermore, to align with the biochemical analysis of liver enzymes, we conducted histopathological analysis to provide further insights into the effects of alcohol and the potential protective role of alpha-asarone. In the ALC group, alcohol-induced liver injury was evident in the histopathological examination. Multiple areas of portal inflammation with evidence of interface hepatitis, an area of confluent necrosis, and scattered foci of lobular inflammation were observed. The presence of portal inflammation, interface hepatitis, and necrosis are characteristic histopathological features associated with alcoholic liver disease (ALD) [27, 28]. These findings validate the association between alcohol consumption and histopathological deterioration in the liver, which has been consistently confirmed in previous studies. The inflammatory infiltrates and necrotic areas observed in the ALC group further emphasize the detrimental effects of chronic alcohol consumption on liver tissue.

In contrast, treatment with alpha-asarone (5, 10 mg/kg) showed noticeable improvements in liver histology. The extent of portal inflammation, interface hepatitis, and necrosis was visibly reduced compared to the ALC group. These histopathological findings align with the biochemical analysis of liver enzymes, suggesting that alpha-asarone may exert protective effects against alcohol-induced liver injury.

Overall, our histopathological analysis provides supportive evidence for the association between alcohol consumption and liver histopathological deterioration. Additionally, it highlights the potential of alpha-asarone to ameliorate alcohol-induced histopathological changes in the liver. Further investigations, including advanced detailed evaluation of liver tissue, are warranted to elucidate the underlying mechanisms and confirm the histopathological benefits of alpha-asarone in ALD.

The inflammatory response to chronic ethanol consumption is characterized by the involvement of proinflammatory cytokines such as interleukin-1 (IL-1), interleukin-6 (IL-6), and tumor necrosis factor-alpha (TNF-α). These cytokines are released by Kupffer cells and peripheral blood monocytes and play critical roles in propagating the inflammatory cascade [29, 30]. The release of these proinflammatory cytokines leads to cell death through necrosis and apoptosis, further accelerating liver damage [31, 32]. Targeting proinflammatory cytokines has shown promise as an effective approach for treating alcoholic liver injury. In our study, we observed a significant increase in the levels of IL-1, IL-6, and TNF-α in the ALC group, indicating liver injury due to alcohol consumption. However, treatment with alpha-asarone (5, 10 mg/kg) mitigated these elevations, suggesting that the expression of proinflammatory cytokines was suppressed which indicates that alpha-asarone has an anti-inflammatory effect. The ability to modulate the expression of IL-1, IL-6, and TNF-α is crucial for the successful treatment of ALD. Our findings are consistent with a study conducted by Tathe et al. [26], which demonstrated the effect of alpha-asarone on IL-1, IL-6, and TNF-α in hepatotoxic rats. They reported similar reductions in the levels of these proinflammatory cytokines with alpha-asarone treatment. Also, Shin et al. [33] investigated the impact of alpha-asarone on memory deficit mice and found that it normalized the levels of IL-1, IL-6, and TNF-α after four days of treatment.

Oxidative stress is considered a major pathological mechanism involved in the initiation and progression of liver injury, including alcoholic liver disease. Various risk factors such as alcohol, drugs, environmental contaminants, and irradiation can induce oxidative stress in the liver, leading to severe liver disorders [34]. Under certain physiological conditions, oxidative stress can be beneficial, as it can enhance biological defense systems through appropriate physical activity and ischemia, as well as trigger apoptosis to prepare the birth canal for delivery [35, 36]. However, in the context of alcohol-induced liver disease, reactive oxygen species (ROS) play a critical role. Ethanol metabolism by alcohol dehydrogenase (ADH) and aldehyde dehydrogenase (ALDH) leads to increased cellular ROS levels, resulting in lipid peroxidation, DNA damage, and liver injury. Moreover, alcohol administration affects the expression of various antioxidant enzymes, including glutathione (GSH), superoxide dismutase (SOD), catalase (CAT), and malondialdehyde (MDA) [37, 38]. In our study, we observed a significant decrease in GSH concentration, SOD activity, and CAT activity in the ALC group compared to the normal group, indicating impaired antioxidant defense mechanisms. However, treatment with alpha-asarone (5, 10 mg/kg) resulted in a significant increase in GSH concentration, SOD activity, and CAT activity compared to the ALC group. Additionally, the ALC group showed a significant increase in MDA content, a marker of lipid peroxidation, compared to the normal group. Conversely, the treatment groups exhibited a marked decrease in MDA content compared to the ALC group. These findings suggest that alpha-asarone can ameliorate the reduction in antioxidant enzyme activities associated with alcohol-induced liver injury and has a potential effect on the treatment of ALD. Consistent with our results, previous studies have also demonstrated the antioxidant activity of alpha-asarone by enhancing GSH, SOD, and CAT in hepatotoxic rats [26]. Similarly, in a study by Sutariya et al. [39] on nephrotic syndrome rats, alpha-asarone significantly restored the depletion of endogenous enzymatic antioxidants.

Autophagy, an intracellular pathway involved in maintaining cellular homeostasis, has been shown to have protective effects against alcohol and drug-induced liver damage [40]. It serves as a mechanism for eliminating damaged organelles and macromolecules through their delivery to lysosomes for degradation. Consequently, autophagy plays a crucial role in preserving cellular homeostasis under both healthy and diseased conditions [41, 42]. Thus, targeting autophagy may offer new therapeutic options for ALD patients. Previous studies have explored the role of AMP-activated protein kinase (AMPK), a marker of autophagy, in ALD [43]. AMPK either directly stimulates autophagy by phosphorylating autophagy-related proteins in the mTORC1, ULK1, and PIK3C3/VPS34 complexes, or indirectly by influencing the expression of autophagy-related genes downstream of transcription factors such as FOXO3, TFEB, and BRD4 [44]. AMPK can also increase autophagic breakdown of mitochondria (mitophagy) by inducing fragmentation of damaged mitochondria in the network and promoting autophagy machinery translocation to damaged mitochondria [44]. Beclin-1 and LC3 are key autophagy-related proteins involved in the autophagy process [45]. Beclin-1 is a molecular platform that assembles an interactome of stimulating and suppressive components that govern the onset of autophagosome formation [46]. During autophagy, autophagosomes engulf cytoplasmic components such as cytosolic proteins and organelles. Concurrently, a cytosolic form of LC3 (LC3-I) is conjugated to phosphatidylethanolamine to generate LC3-phosphatidylethanolamine conjugate (LC3-II), which is recruited to autophagosomal membranes [47].

Alcohol consumption has been shown to decrease intracellular AMPK levels, while increasing the levels of LC3 and Beclin-1, thereby enhancing autophagic flux [48, 49]. In our study, we observed a significant decrease in AMPK expression in the ALC group compared to the normal group. Conversely, treatment with alpha-asarone significantly increased AMPK expression in the treated groups. Additionally, the ALC group showed a significant increase in Beclin-1 and LC3 expression compared to the normal group, whereas the treatment groups exhibited a marked decrease in Beclin-1 and LC3 expression. Our results suggest that alpha-asarone has a potential effect in the treatment of ALD via modulating AMPK, Beclin-1, and LC-3 activity. These findings align with those of Das et al [50] who reported that alpha-asarone enhanced AMPK expression in a diabetic-hepatocellular carcinoma condition. Furthermore, alpha-asarone administration was found to inhibit the expression of LC3 in rats with cerebral ischemia-reperfusion stroke [51].

Apoptosis, also known as programmed cell death or cellular suicide, is a fundamental process involved in the elimination of damaged or unwanted cells. Dysregulation of the apoptotic pathway plays a significant role in the development and progression of liver diseases, including ALD [52]. The tumor suppressor protein p53 has been implicated in the pathogenesis of liver disease, and its activation is crucial in ethanol-induced hepatocyte apoptosis, as demonstrated by the protective effect of p53 genetic ablation against ethanol-induced liver damage [53, 54]. In our study, immunohistochemical analysis using an anti-p53 antibody was conducted to evaluate p53 expression as an apoptosis marker. The results revealed strong nuclear and cytoplasmic expression of p53 in hepatocytes of the ALC group compared to the normal group. However, treatment with alpha-asarone in the treated groups resulted in a significant reduction in cytoplasmic p53 expression in hepatocytes. These findings are consistent with a study by Shi et al. [55] where both alpha and beta asarone reduced p53 expression in an Aβ-induced inflammatory response in the PC12 cell model. All these findings provide significant evidence that alpha-asarone may be a novel bioactive agent for the treatment of alcoholic liver disease.

## 5. Conclusions

Collectively, these findings provide compelling evidence that alpha-asarone may serve as a promising bioactive agent for the treatment of alcoholic liver disease. Its beneficial effects are attributed to its ability to attenuate the inflammatory response by suppressing proinflammatory cytokines such as IL-1, IL-6, and TNF-α, as well as its antioxidant properties, which enhance the activities of GSH, SOD, and CAT while reducing lipid peroxidation. Furthermore, alpha-asarone exhibits modulatory effects on autophagy markers, such as AMPK, Beclin-1, and LC3, and it can mitigate apoptosis by reducing p53 expression. Further investigations are warranted to elucidate the underlying molecular mechanisms and evaluate the therapeutic potential of alpha-asarone in clinical settings.

## Abbreviations

ALP: Alkaline Phosphatase
AST: Aspartate Aminotransferase ALT: Alanine Aminotransferase MDA: Malondialdehyde
GSH: Reduced Glutathione CAT: Catalase
SOD: Superoxide Dismutase
TNFα: Tumor Necrosis Factor Alpha IL-1: Interleukin-1
IL-6: Interleukin-6
ELISA: Enzyme-Linked Immunoassay
AMPK: Adenosine Monophosphate-Activated Protein Kinase
LC3: Microtubule-Associated Protein Light Chain 3
NADPH: Nicotinamide Adenine Dinucleotide Phosphate
CYP2E1: Cytochrome P450 2e1
ROS: Reactive Oxygen Species
DNA: Deoxyribonucleic Acid
ALD: Alcoholic Liver Disease
CNS: Central Nervous System
CVS: Cardiovascular System ETOH: Ethanol
NIAAA model: Chronic Ethanol Feeding Plus a Single Binge.
IHC: Immunohistochemistry
H&E: Hematoxylin and Eosin Pg: Picogram
ADH: Alcohol Dehydrogenase ALDH: Aldehyde Dehydrogenase

## Acknowledgments

Funding: This research did not receive any specific grant from funding agencies in the public, commercial, or not-for-profit sectors.

## Credit authorship contribution statement

Conceptualization of the research idea, data curation, methodology development and final revision were implemented by N.M.E; Literature review, experiments, data collection, data analysis, editing and interpretation were implemented by N.A.M.A; Methodology development, data analysis, editing, interpretation and revision were implemented by A.M.A.; Literature review, data analysis, and revision were implemented by A.R.

